# PlantScience.ai: An LLM-Powered Virtual Scientist for Plant Science

**DOI:** 10.1101/2025.10.24.684337

**Authors:** Haopeng Yu, Shasha Zhou, Mingyu Huang, Ling Ding, Yuxuan Chen, Yinru Wang, Yingyu Ren, Nuo Cheng, Xinya Wang, Jie Liang, The John Innes Centre and The Sainsbury Laboratory Collaboration, Huakun Zhang, Yiliang Ding, Ke Li

**Affiliations:** Department of Cell and Developmental Biology, John Innes Centre, Norwich Research Park, Norwich, NR4 7UH, UK; Department of Computer Science, University of Exeter, Exeter, EX4 4RN, UK; Key Laboratory of Molecular Epigenetics of the Ministry of Education, Northeast Normal University, Changchun, 130024, China; James Watt School of Engineering, University of Glasgow, Glasgow, G12 8QQ, UK

## Abstract

The accelerating growth of plant science literature presents a major challenge for researchers seeking to extract accurate, up-to-date knowledge from an increasingly fragmented and domain-specific corpus. General-purpose large language models (LLMs), while powerful, often misinterpret plant science terminology and lack mechanisms for source traceability. We created *PlantScience*.*ai*, a virtual plant biology scientist powered by our automated scientific knowledge graph construction pipeline (AutoSKG). *PlantScience*.*ai* exhibits expert-level reasoning in plant biology and maintains scholarly rigour in its citations. Through continuous learning, it integrates the latest research, ensuring that its knowledge base remains current and scientifically robust. Apart from providing the answers to the scientific questions, *PlantScience*.*ai* can interact with human scientists, follow instructions, and retrieve information with citation awareness, grounding each response in primary sources to ensure accuracy and verifiability. *PlantScience*.*ai* marks a pivotal advance toward a collaborative scientific paradigm in which virtual and human plant scientists work synergistically to accelerate discovery while preserving the unique value of human insight. *PlantScience*.*ai* is available at https://plantscience.ai.

## Main

Over the past century, plant biology has undergone a profound transformation, driving advances in agricultural innovation, ecosystem stewardship and fundamental biological understanding (Smith et al., 2009). Whilst concurrent breakthroughs in molecular genetics, developmental biology, environmental physiology, and ecological modelling have generated vast amounts of knowledge, documented in published literature and various specialised databases, the rapid expansion of information has led researchers to seek more efficient strategies for extracting relevant content from this wealth of knowledge. Although large language models (LLMs) offer promising capabilities for information synthesis, their application in biology still faces significant limitations (Zhang et al., 2025a). LLMs are prone to fabricating plausible but erroneous citations and statements, or ‘hallucinations’, a shortcoming that is exacerbated in specialised domains, where deep, field-specific expertise is indispensable (Farquhar et al., 2024). Furthermore, compared to general knowledge, plant science contains numerous technical terms and abbreviations that often confuse general-purpose LLMs. For instance, in plant science, “ABA” refers to abscisic acid, whereas in general knowledge contexts, it more commonly stands for applied behaviour analysis. Similarly, asking “Is FT the main target of CO?” may confuse most LLMs. The question actually refers to whether the FLOWERING LOCUS T gene is a target of the CONSTANS transcription factor. Such semantic ambiguity, coupled with the opacity of commercial LLMs and their lack of traceable information sources, further limits their scientific utility. To address this gap, we posit that integrating domain-specific knowledge into generative AI systems, implemented with transparency and verifiability, is a crucial step towards developing virtual scientific assistants capable of meaningful collaboration with human researchers.

Here, we present *PlantScience*.*ai*, a virtual plant scientist that serves as an intelligent research companion. It possesses extensive knowledge of plant biology and continuously expands its understanding by learning from new research, whilst engaging with plant biologists through an intuitive interface. Through natural language conversation, researchers can query complex biological questions and receive evidence-based answers with citation transparency. Beyond the information retrieval, *PlantScience*.*ai* synthesises knowledge across sources, visualises interconnected concepts through interactive knowledge graphs, and enables researchers to verify and explore the evidential basis of every response.

We trained *PlantScience*.*ai* by emulating the process of educating human plant scientists. Like students who first master fundamental biological principles such as genes, proteins, and pathways, *PlantScience*.*ai* is trained on core biological concepts to build foundational understanding. It further learns to integrate how different studies describe the same concepts, synthesising diverse perspectives into a coherent understanding. For example, when multiple studies assign different functions to a single gene, *PlantScience*.*ai* integrates these insights to produce a unified interpretation. Moreover, it continuously refines its knowledge base, reassessing existing information in light of new research, to ensure its understanding evolves alongside scientific progress.

To implement this learning paradigm, we developed an automated scientific knowledge graph construction pipeline (AutoSKG) that enables *PlantScience*.*ai* to acquire, organise, and continuously update its knowledge (Figure 1A). In AutoSKG, all knowledge is represented as a heterogeneous network containing entities and relationships, forming a knowledge graph (KG). We first predefined knowledge categories specific to plant science, ranging from genes, proteins, and metabolites to biological pathways and experimental toolsets. The pipeline dynamically expands these categories when novel entity types are identified during learning, currently encompassing approximately 1,000 distinct knowledge categories. To enable fully automated knowledge extraction, we employed LLM-based named entity recognition (NER) and text summarisation techniques to identify biological entity names and their relationships in the raw data and generate descriptions for them by synthesising information from different sources (Al-Moslmi et al., 2020; Edge et al., 2025; Zhang et al., 2025b). All entity and relationship descriptions can be linked back to the original source materials, enabling researchers to trace and verify the underlying data. When new research emerges, AutoSKG employs an incremental update mechanism that re-synthesises knowledge by integrating novel findings with existing information, thereby enabling continuous learning. Through the AutoSKG pipeline, we construct a knowledge graph database, the plant science knowledge graph (PSKG). PSKG is a dynamic, domain-specific knowledge base that serves as the core of *PlantScience*.*ai*, encompassing comprehensive plant science knowledge drawn from diverse sources and continuously updated through automated literature acquisition and user contributions. In June 2025, our PSKG contained approximately 1.2 million biological entities and 4.0 million edges representing their interconnections. By September 2025, continuous learning through AutoSKG had expanded PSKG to 1.8 million entities and 6.5 million edges. We further utilised our AutoSKG to expand the knowledge base by integrating the PlantConnectome, a recently published LLM database derived from approximately 71,000 research articles (Lim et al., 2025). The current (Oct 2025) PSKG now comprises 4.0 million entities and 10.8 million edges, numbers that continue to increase as new literature is processed.

**Figure 1.**
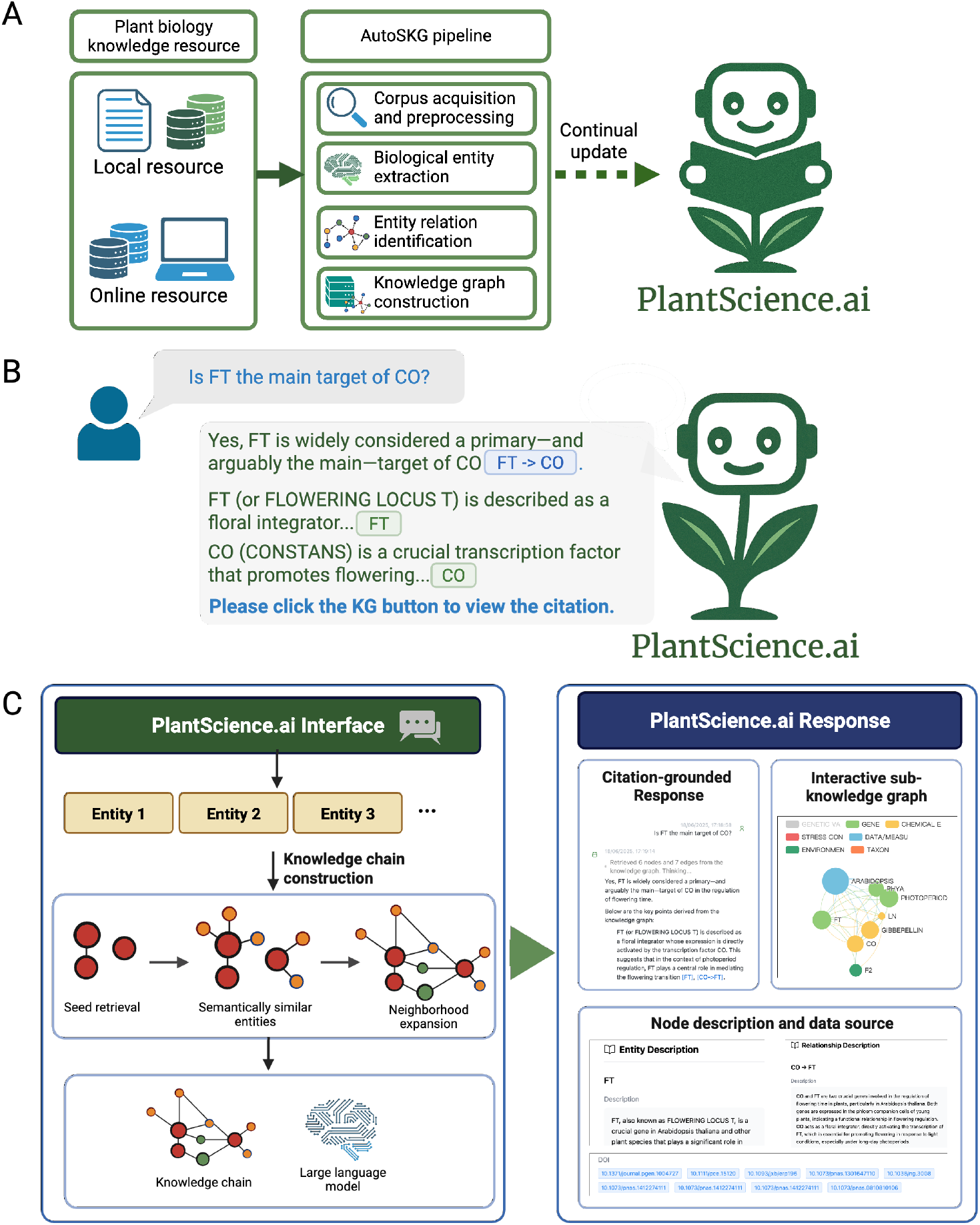
Schematic overview of *PlantScience*.*ai*. **(A) AutoSKG pipeline for knowledge graph construction**. Plant biology literature from local and online sources is processed through corpus acquisition, entity extraction, relation identification, and graph construction, enabling continual updates of the plant science knowledge graph (PSKG). The robot holding a book symbolises PlantScience.ai during the knowledge acquisition and updating phase, where it learns and expands the Plant Science Knowledge Graph (PSKG) through the AutoSKG pipeline. **(B) The example of the citation-grounded response**. In response to user queries, *PlantScience*.*ai* retrieves relevant information and generates natural language responses with traceable citations displayed as KG buttons. These buttons can be clicked to access detailed information about specific entities or relationships. The robot without a book represents PlantScience.ai in the interactive phase, where it engages with users and generates responses based on the knowledge it has previously learned. **(C) Knowledge chain construction and response interface**. Entities in the query are used to construct a semantic knowledge chain. LLM integrates to produce responses, alongside interactive graphs and structured node-level descriptions.

Yet acquiring knowledge is only half the challenge, *PlantScience*.*ai* is also designed to learn to retrieve the right information and communicate it effectively, much as a trained scientist learns to navigate literature and construct evidence-based arguments. To achieve this, we implemented graph-based retrieval-augmented generation (RAG), enabling PlantScience.ai to locate relevant knowledge and trace logical connections between concepts when answering queries (Figure 1C). Once a query is submitted, *PlantScience*.*ai* first employs NER to identify key entities within the user’s question (Figure 1C) (Al-Moslmi et al., 2020). These entities are then processed using graph-based RAG technology to retrieve the most relevant entities from the KG (Edge et al., 2025). Subsequently, we perform graph traversal from the target nodes and semantically similar nodes within the KG, leveraging LLM-based reasoning and semantic-similarity strategies to identify additional relevant edges and nodes. This process constructs a “knowledge chain” composed of intermediate entities and relationships that connect the relevant query. Moreover, the factuality and relevance of the responses generated by *PlantScience*.*ai* can be traced back to its corresponding entities, relationships, and data sources, allowing users to verify the factual basis of the information directly. Users have the option to visualise the relevant sub-KG through the web interface (Figure 1C). Such visualisation enables interactive exploration of the entities and relationships, along with their detailed descriptions and links to the original source data, fostering a deeper understanding and discovery. Meanwhile, each time it responds, PlantScience.ai will search online for updated knowledge to improve its answer, and these updates will subsequently be integrated into PSKG through the AutoSKG pipeline. During the conversation, we employ client-side storage, ensuring that all chat history is archived exclusively on the client device to maintain data privacy and security.

The *PlantScience*.*ai* underwent over a year of internal testing, with iterative improvements guided by feedback from domain experts. The trial engaged about 150 scientists from the John Innes Centre (JIC) and the Sainsbury Laboratory (TSL), contributing to system refinement through internal testing and expert consultation, helping identify knowledge gaps, improve response accuracy, and enhance handling of domain-specific terminology. To evaluate scientific accuracy and depth, we designed a domain-expert evaluation to benchmark its responses to domain-specific questions against those generated by several leading LLMs. The responses were assessed according to four key criteria: factual accuracy, relevance, and comprehensiveness. Benchmark results demonstrated that *PlantScience*.*ai* outperformed the other chatbots in these dimensions (Methods, Figure S1). These findings indicate that *PlantScience*.*ai* is substantially more reliable than general-purpose chatbots when addressing queries within the plant science domain.

*PlantScience*.*ai*, the pioneering virtual scientist for plant biology, combines a general-purpose LLMs with a rigorously curated, domain-specific KG to deliver evidence-anchored, verifiable answers. While plant-specific LLM agents such as PlantGTP, mainly developed for *Arabidopsis thaliana*, and SeedLLM, designed for rice, have emerged based on static knowledge databases, *PlantScience*.*ai* supports continuous knowledge integration across a broad spectrum of plant science (Zhang et al., 2025c; Yang et al., 2025). Furthermore, aside from continuous learning capabilities, PlantScience.ai ensures source-traceable conversations with transparency, implements client-side storage for data privacy, and features dynamically displayed knowledge graphs for intuitive concept exploration. The platform incorporates a robust feedback system and has demonstrated stability through extended real-world usage. Although built for plant science, its architecture, which includes a PSKG with continuous learning capabilities enabled by the AutoSKG pipeline, a graph-based RAG, and an intuitive interface, offers a transferable blueprint for disciplines that struggle with information overload. Users may also leverage our open-source AutoSKG pipeline to construct private, personalised knowledge graphs from their own unpublished or sensitive documents.

To exemplify the potential of human-AI synergy in scientific research, we created *PlantScience*.*ai*, a virtual scientist tailored to plant biology that demonstrates how LLM, when coupled with curated domain knowledge. *PlantScience*.*ai* is committed to an open and collaborative community where we invite global researchers to engage in the further development, evaluation, and benchmarking of virtual scientists, finally leading to a new human-AI synergistic scientific discovery paradigm.

## Data Availability

The source code for AutoSKG is available at the GitHub repository: https://github.com/COLA-Laboratory/autoSKG.

## Acknowledgements

This research was supported in part by the NBI Computing infrastructure for Science (CiS) group and Informatics platform. We would like to thank the John Innes Centre and the Sainsbury Laboratory consortium for their participation in the *PlantScience*.*ai* testing.

## Author information

### Author notes

Haopeng Yu, Shasha Zhou, Mingyu Huang and Ling Ding contributed equally to this work.

### Authors and Affiliations

Department of Computer Science, University of Exeter, Exeter, EX4 4RN, UK. Shasha Zhou, Mingyu Huang, Yuxuan Chen, Ke Li

Department of Cell and Developmental Biology, John Innes Centre, Norwich Research Park, Norwich, NR4 7UH, UK

Haopeng Yu, Yiliang Ding

Key Laboratory of Molecular Epigenetics of the Ministry of Education, Northeast Normal University, Changchun, 130024, China

Ling Ding, Yinru Wang, Yingyu Ren, Nuo Cheng, Xinya Wang, Jie Liang, Huakun Zhang

## Contributions

K.L., Y.D. and H.Z. conceived the study; H.Y., S.Z., M.H., L.D., H.Z., K.L., and Y.D. designed the study; H.Y. implemented the full-stack development of PlantScience.ai; H.Y., L.D., Y.W., Y.R., N.C., X.W., and J.L. collected the data; S.Z., M.H., and Y.C. developed the knowledge graph; H.Y., S.Z., M.H., and Y.C. developed AutoSKG; K.L., H.Z., and Y.D. supervised the process and analyses; H.Y., S.Z., M.H., K.L., H.Z., and

Y.D. wrote the manuscript with input from all authors; The John Innes Centre (JIC) and the Sainsbury Laboratory (TSL) collaboration participated in one year of evaluation and feedback for PlantScience.ai.

## Corresponding authors

Correspondence to <u>Ke Li</u>, <u>Huakun Zhang</u> and <u>Yiliang Ding</u>.

## Ethics declarations

Ethics approval and consent to participate Not applicable.

## Supplemental figure

**Figure S1.**
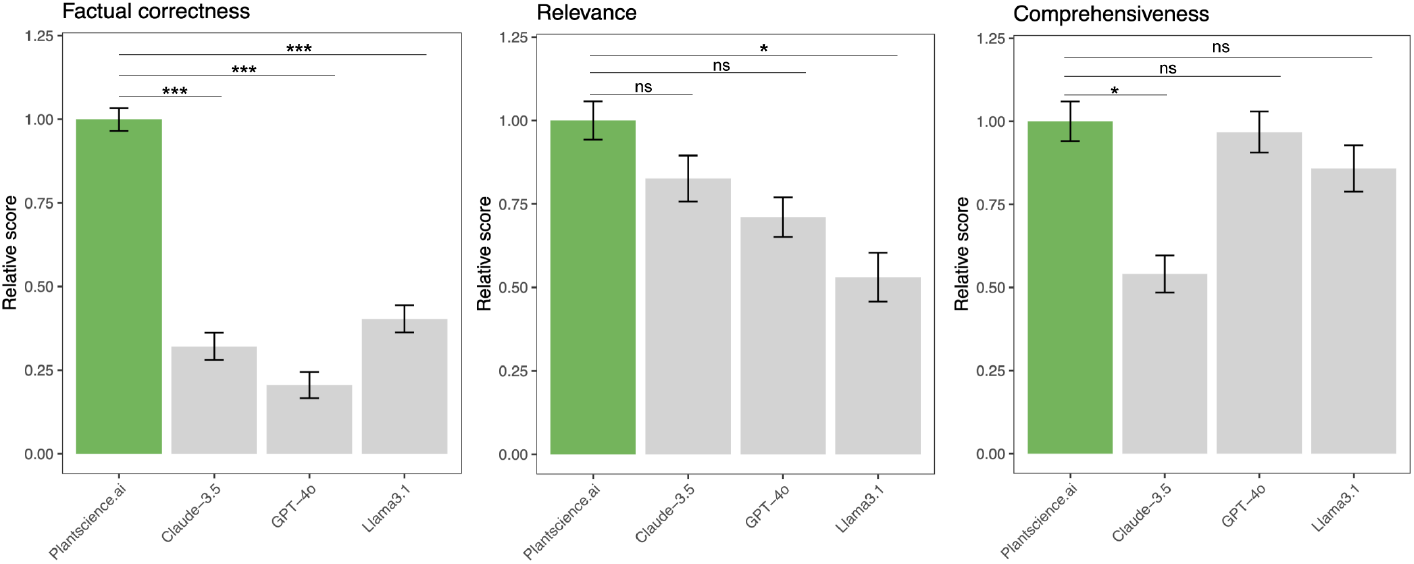
Benchmark of PlantScience.ai against leading LLMs. Benchmark comparison on factual correctness, relevance, and comprehensiveness. Bars indicate mean normalised scores across evaluation queries, and error bars represent the standard error of the mean (SEM). Statistical significance was determined using t-tests: P < 0.05, P < 0.01, P < 0.001c.

## Supplemental information (SI)

### Methods

#### AutoSKG pipeline

Data processing. The AutoSKG pipeline starts from preprocessing all the collected data. Specifically, we parsed all non-textual (e.g., PDF) files using optical character recognition (OCR) technique to obtain raw text corpus. We then assigned a unique hashing ID for each file to enable later backtracking. Next, for all textual data, we applied a preprocessing pipeline which involves standardising encoding schemes, correcting special characters as well as filtering out line breaks. We converted all text to lowercase to enhance processing efficiency by LLMs.

Entity and relationship extraction. To extract biological entities and relationships, we prompted an LLM to go through the entire text corpus. For this purpose, we used GPT-4o-mini to keep a nice balance between performance and cost. Each time, the model is provided with an extraction of the text with 1,500-token in length and 200-token overlap between consecutive windows. The model is instructed to identify any biological entities and quote the context in which these entities appear, thus essentially performing the NER task. To enhance extraction quality and avoid hallucinations or missing entities, we applied in-context learning by incorporating five expert-crafted examples in the prompt. We also invoked multiple rounds of scrutiny with follow-up critical analysis questions (e.g., “does the current extraction cover all biological entities appear in the original text?”, “does the current extraction contain any biological entity that actually do not exist in the original text?”) to ensure full coverage while avoiding hallucinations. After extracting all entities from the entire text corpus, we conducted another round of LLM scrutiny to identify any explicitly stated biological relationships between the previously extracted entities using a similar prompting method above and recorded the context involving such statements.

Once all the entities and relationships are extracted, we mapped all the results into a structured knowledge graph, wherein nodes are extracted biological entities, and edges indicate stated relationships between them. During this process, synonyms and spellings were standardised and merged to avoid redundancy. We also generated a final description for each node and entity by summarising all the relevant text pieces describing them collected during the extraction process. For each node, we additionally prompted GPT-4o to assign a category for it from a predefined list, which includes. During the PlantConnectome integration, novel entities and relationships were directly incorporated into the knowledge graph, whilst pre-existing entities and relationships were reconciled using AutoSKG's summarisation pipeline to integrate complementary information from both sources.

#### Construction of Plant Science Knowledge Graph

The Plant Science Knowledge Graph (PSKG) is constructed and continuously maintained through our automated AutoSKG pipeline. The initial version of the PSKG focused on model organisms such as *Arabidopsis* and key crops including wheat, rice, and maize, supplemented with structured, community-contributed data from researchers at the John Innes Centre (JIC) and The Sainsbury Laboratory (TSL). These initial datasets were manually curated by plant experts at JIC/TSL prior to processing by AutoSKG to ensure a high-quality foundation.

The PSKG has since been substantially expanded to incorporate tens of thousands of full-text articles from diverse peer-reviewed journals across plant science. We have continuously broadened the data sources to encompass literature from a wider phylogenetic range. Furthermore, the PSKG integrates the PlantConnectome knowledge graph, enriching the knowledge base with additional structured biological relationships.

The PSKG remains current through a hybrid two-mode update that enables continuous learning and knowledge expansion. First, when users submit queries, the system performs web searches on the same topic to identify recent publications that have not yet been integrated into the PSKG. These newly discovered studies are automatically processed through the AutoSKG pipeline and incorporated into the knowledge graph. Second, users can directly upload their own research findings or domain-specific knowledge via (https://plantscience.ai/upload ), which are similarly integrated into the PSKG via AutoSKG. To ensure computational efficiency, all newly acquired publications from web searches and user-submitted data are processed in collective batch updates that run weekly.

#### Graph-based RAG

For each user input, we first use a GPT-4o model to perform NER on it to identify key biological concepts involved. We then search within the KG for relevant nodes. To do so, we create embedding vectors for all nodes in the KG and the identified biological concepts in a latent semantic space, in which semantically similar concepts would lie close to each other. Here, we use the voyage-3-large model from Voyage AI, which demonstrates state-of-the-art performance across a wide range of text embedding benchmarks, outperforming comparable models such as OpenAI-v3-large by approximately 9.7% on average (Voyage AI, 2025). After constructing the embedding vectors, we search for nodes in the KG whose embeddings yield the highest cosine similarities with the key concepts in the user input.

We then construct the “knowledge chain” through a graph-based RAG approach involving multi-stage subgraph retrieval and expansion (Edge et al., 2025). First, seed entities identified through NER are retrieved from the PSKG along with their immediate neighbours, forming the initial query-relevant subgraph. Second, we leverage embedding-based semantic similarity to identify and retrieve semantically similar entities, effectively capturing conceptually related nodes that may not be directly connected in the graph topology. Third, we perform neighbourhood expansion by traversing the k-hop neighbourhoods of both the seed and semantically similar entities, applying Dijkstra's algorithm to enumerate the shortest paths between each pair of relevant nodes (Fan and Shi, 2010). This multi-stage graph RAG pipeline ensures comprehensive retrieval of both explicit structural connections and implicit semantic associations within the knowledge graph. Each resulting knowledge chain comprises a sequence of intermediate entities and typed semantic relationships that establish the logical connections between relevant biological concepts. The aggregated textual descriptions and metadata associated with each entity and edge in the knowledge chain provide comprehensive provenance information. We then feed this enriched contextual knowledge to the LLM, enabling it to generate evidence-grounded, verifiable responses that directly address user queries whilst maintaining full traceability to the underlying data sources.

#### Full-stack development of PlantScience.ai

The web interface of PlantScience.ai was developed as a comprehensive web-based application utilising modern frontend technologies to provide an intelligent research assistant for plant science. The front-end architecture employs “Vue.js 3” with the composition API (vuejs.org). For frontend-backend communication, we integrated “Axios” as the HTTP client (axios-http.com), enabling real-time streaming responses and continuous parsing of incoming data, critical for progressively delivering PlantScience.ai's answers as they are generated. The chat interface supports context-aware conversations, with responses rendered via a “Markdown rendering stack” that allows rich formatting of scientific notation, citations, and structured information. Interactive knowledge graph visualisations are powered by ECharts (echarts.apache.org), enabling users to dynamically explore biological relationships. To prioritise user privacy, we employed Pinia (pinia.vuejs.org) for state management, which stores all conversation history and user data exclusively on the client side rather than transmitting it to servers. The application implements internationalisation support for multiple languages and incorporates user authentication systems with secure token-based authorisation. The development methodology adheres to modern web standards, incorporating progressive web application capabilities to ensure cross-platform compatibility across desktop and mobile devices.

The server-side of PlantScience.ai was developed as a robust API server utilising modern backend technologies. The system architecture uses “FastAPI” (fastapi.tiangolo.com) as the primary framework, leveraging its asynchronous request handling to deliver efficient, real-time streaming responses. Knowledge extracted by AutoSKG is stored in a “Neo4j graph database”, which supports comprehensive CRUD operations (create, read, update, delete) for nodes and relationships, enabling automated knowledge updates and maintenance. To facilitate intelligent information retrieval, we implemented a hybrid search system that combines exact, fuzzy, and semantic matching. We also embedded intent detection algorithms to determine whether queries require knowledge graph interrogation. Session management handles conversation data storage, automatic expiry, and temporary caching of graph data to optimise performance. The authentication system utilises JWT tokens with bcrypt password hashing, supporting user registration with email verification, login authentication, password reset functionality, and usage limits (chat quotas and feedback counts). An integrated email service manages verification emails, password resets, and feedback notifications. Performance optimisations include batch querying, data compression (removing large fields such as embeddings), timeout controls, and entity indexing to ensure a responsive user experience even with complex queries.

#### Benchmark Design and Evaluation

To assess PlantScience.ai's performance against existing general-purpose LLMs, we developed a benchmark comprising 60 questions spanning molecular biology, genetics, physiology, and pathology in plant science (Table S1). Questions were curated and validated by senior researchers from the John Innes Centre and The Sainsbury Laboratory to ensure scientific relevance and real-world applicability (Supplementary Table 1). Four models were evaluated: PlantScience.ai, Claude 3.5 Sonnet (Anthropic), GPT-4o (OpenAI), and Llama 3.1 8B (Meta). All models were queried under identical conditions using default parameters without additional prompt engineering. To validate scientific accuracy and depth, we designed a domain-expert evaluation. Model responses were evaluated using a standardised rubric based on three metrics, each scored on a 1–10 scale: Factual Accuracy (scientific correctness verifiable against primary literature), Relevance (direct and complete addressing of the question), and Comprehensiveness (depth of information, with mechanistic and contextual details). To ensure objectivity, all responses were anonymised and randomly ordered before evaluation. Next, for the same question and the same evaluator, the four scores for the four models are normalised to a 0-1 scale to ensure a fair comparison.

